# Breast cancer metabolism and responsiveness to dichloroacetate: relationships with ^15^N and ^13^C natural abundance

**DOI:** 10.64898/2026.03.09.710495

**Authors:** Illa Tea, Marine P. M. Letertre, Julien Boccard, Anne-Marie Schiphorst, Sophie Blanchet, Mikael Croyal, Anneke C. Blackburn, Guillaume Tcherkez

## Abstract

**Background:** Metabolic reprogramming is a hallmark of breast cancer (BrCa), with alterations in glycolysis, glutamine metabolism, and the urea cycle contributing to tumour progression. Dichloroacetate (DCA), a pyruvate dehydrogenase kinase (PDK) inhibitor, shifts metabolism toward oxidative phosphorylation and has been proposed as a therapeutic agent. While isotope tracing is well-established, natural isotope abundance (δ¹³C, δ¹⁵N) is emerging as a biomarker of metabolic alterations in cancer.

**Methods:** We investigated the relationship between isotope composition and metabolism in BrCa using two BALB/c mouse mammary tumour models (V14 and 4T1) and assessed the effects of DCA treatment using metabolomics, lipidomics and isotopomics.

**Results:** V14 and 4T1 tumours exhibited isotopic patterns similar to human tumours, with δ¹³C enrichment and δ¹⁵N depletion relative to non-cancerous mammary tissue. V14 tumours were more δ¹⁵N-depleted than 4T1, reflecting differences in nitrogen metabolism. Multivariate analysis integrating isotopic, metabolomic, and lipidomic data revealed isotopic features as key discriminators between tumours and normal tissues. Compared to V14, 4T1 tumours were enriched in TCA intermediates, sphingolipids, and amino acids, whereas V14 tumours showed elevated glutaminolytic and nitrogenous metabolites. DCA treatment differentially affected tumour growth, with V14 tumours more sensitive than 4T1. DCA altered nitrogen metabolism, increasing the arginine-to-ornithine ratio, and modulating δ¹⁵N values in a tumour-specific manner increasing V14 and decreasing 4T1 δ¹⁵N values. DCA had little effect on δ¹³C. δ¹³C values were primarily determined by the balance between lipid and TCA cycle metabolites, rather than glycolytic flux. δ¹⁵N variation was linked to nitrogen metabolism, including urea cycle intermediates and sphingolipid composition, with a potential role for choline-related fractionation in δ¹⁵N depletion. Altered gene expression of *Hacd2* and *Acot12* in V14 tumours after DCA treatment was reflected in shorter fatty acid tails in phosphatidyl cholines, supporting the lipidomics data.

**Conclusions:** These findings support the hypothesis that cancer-associated metabolic reprogramming influences natural isotope abundance. Correlations between isotope shifts and metabolic signatures highlight the potential of lipid-derived δ¹⁵N as a biomarker of tumour metabolic state, with implications for noninvasive metabolic profiling in BrCa.

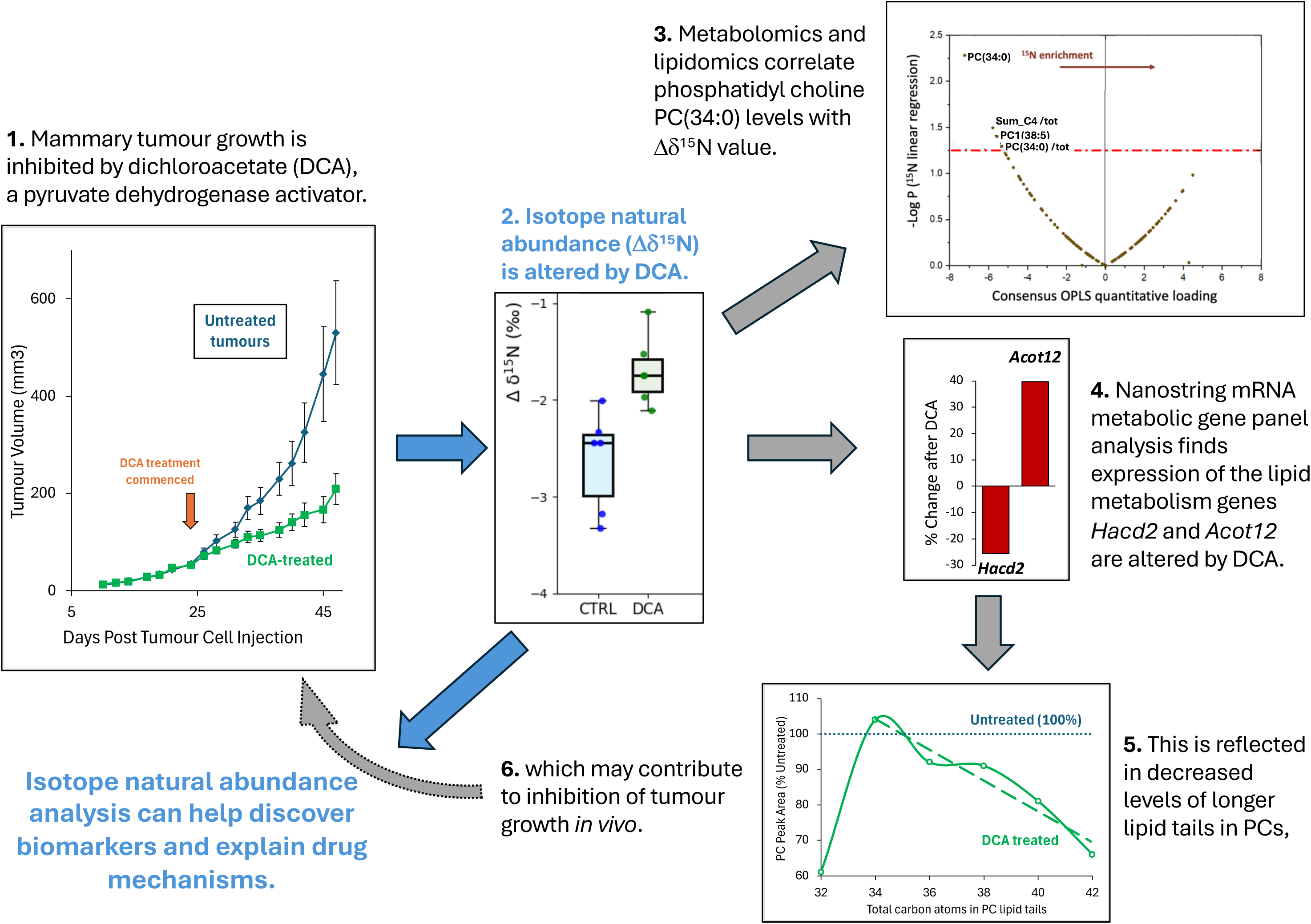

## Background

Intense efforts are currently being devoted to improving treatments and early diagnosis of breast cancer (BrCa), and identifying reliable biomarkers to monitor treatment response. Altered metabolism has been recognised as a hallmark of cancer [1, 2] and thus identifying biomarkers and targeting cancer cell metabolic reactions (e.g. with inhibitors) are viewed as promising strategies for cancer monitoring and therapy. Key metabolic events observed in many cancer cells include elevated glucose uptake and glycolysis, lactate production, increased glutamine metabolism, deregulated urea cycle [3], and altered fatty acid oxidation and synthesis [4, 5].

Amongst clinical studies with metabolic inhibitors, the use of glycolysis inhibitors has provided encouraging data [6, 7]. Inhibitors of pyruvate dehydrogenase kinase (PDK) have some potential to control glycolysis in cancer cells. In fact, PDK inhibition prevents phosphorylation of the pyruvate dehydrogenase (PDH) complex and thus its down-regulation. PDH, when dephosphorylated and active, oxidises pyruvate to CO_2_ and acetyl-CoA, fuelling the tricarboxylic acid (TCA) cycle, and thus redirecting pyruvate from lactate synthesis to catabolism [8]. Also, a relationship has been demonstrated between PDK activity and metastatic potential in breast cancer [9]. Dichloroacetate (DCA) is a well-known PDK inhibitor that has been shown to be effective in reducing the proliferation of various cancer cell lines [10–16]. In effect, DCA decreases aerobic glycolysis, leading to a reduction in lactate production and a stimulation of mitochondrial metabolism, thereby antagonising the suppression of mitochondrion-dependent apoptosis [17, 18]. DCA also leads to a decrease in cytosolic concentration of glycolytic intermediates consumed by the oxidative pentose phosphate pathway [17], contributing to its anti-proliferative effects. In addition to direct metabolic effects, DCA has been found to decrease HIF-1α activity and increase p53 activity, inhibiting glucose uptake, tumour angiogenesis, tumour perfusion, and tumour growth [17, 19, 20]. DCA has been tested in a number of clinical trials (often in combination with other treatments), with some promising results [21–24].

Alongside metabolism manipulation with inhibitors such as DCA, the identification of metabolic signatures of cancer using metabolomics and lipidomics has emerged as a crucial tool for cancer biology. These ‘omics’ technologies can help to identify metabolic pathways and thereby design new metabolic targets for cancer therapy. Many studies have tried to explore BrCa biomarkers using metabolomics (for a recent review, see [25]). Despite some variability between patient cohorts, dietary habits etc., metabolic biomarkers found in tumours via metabolomics are generally consistent with the well-known increase in glycolysis, glutamine utilisation and the onset of the urea cycle in cancer cells (for recent examples, see [26, 27]). Surprisingly, only a few studies have been conducted to evaluate the specific metabolic effects of DCA [28–30]. In the case of mouse lung cancer, DCA has been shown to cause an increase in citrate content and a decrease in nucleotide precursors in the serum, reflecting the enhancement of the TCA cycle and the downregulation of base synthesis [30].

Direct information on metabolism can also be gained from stable isotopes. Typically, the isotope abundance is helpful to interpret changes in metabolite contents, which can be impacted by multiple reactions (metabolic crossroads) or concurrent changes in biosynthesis or consumption. Breast cancer cell metabolism has often been followed using isotope tracing (after isotopic labelling) [9, 31–37]. Taken as a whole, depending on glucose and glutamine availability and cell lines, cancer cells use glycolysis at a high rate to produce lactate and oxaloacetate via anaplerosis (pyruvate carboxylase), or consume glutamine, which can be converted back to citrate and oxaloacetate via ATP citrate lyase, thereby feeding glucogenesis (for a recent study, see [38]). Also, it has been shown that serine metabolism is crucial to cancer cells to feed C_1_ metabolism [39]. ^13^C-tracer studies have shown that DCA leads to a decrease in the glycolytic flux and the pentose phosphate pathway, and increases glutamine metabolism in lactate-producing tumour cells [40]. In pancreatic cancer cells, DCA causes a strong reduction of ^13^C incorporation from glucose to amino acids, mostly due to the reduction in the glycolytic flux [28].

While isotope tracing is useful, it has some limitations. First, regulations may prohibit the use of stable isotopes when ethics approval is not granted. Second, this method usually disregards differences in reaction rates between isotopic forms (referred to as isotope effects). Although isotope effects are rather small (a few ‰) with ^13^C or ^15^N, variations in isotope abundance (referred to as isotope composition or δ-value) are easily measurable with isotope ratio mass spectrometry (IRMS). In principle, changes in metabolism (and thus metabolic isotope effects) should be associated with changes in δ^13^C and δ^15^N, offering some potential to identify isotopic biomarkers of cancer cell metabolism. That is, it is possible to exploit differences in natural isotope content between samples (typically cancer vs. non-cancer cells), without the need to carry out isotope labelling.

Recent advances in analytical methods have facilitated the use of natural stable isotope abundance in biomedicine [41–43]. So far, isotope natural abundance has been used to study normal metabolism of healthy individuals [44–48] and various diseases implying metabolic modifications [49, 50], including cancer [51–60]. In particular, it has recently been shown that cancer cell lines cultured in vitro are naturally ^15^N-depleted and ^13^C-enriched compared to non-cancer cells; similarly, tumour tissues collected from patients are naturally ^15^N-depleted and ^13^C-enriched compared to non-cancer adjacent tissues [54]. This change in isotope composition essentially relates to the lower content in (naturally ^13^C-depleted) lipids, higher content in (naturally ^13^C-enriched) organic acids, and enhanced N metabolism to recycle ammonia liberated by glutamine consumption in cancer cells. Cancer cells contain more (naturally ^15^N- depleted) arginine, leading to a general ^15^N-depletion in cellular nitrogen [43, 54]. Also, the higher glycolytic flux leads to a decline in the overall isotope fractionation against ^13^C, leading to relatively ^13^C-enriched lactate.

However, it remains uncertain as to whether ^13^C and ^15^N natural abundance is also impacted by variations in cancer cell metabolome, differences in cancer mutation profile, or by metabolic inhibitors such as DCA. These questions are of importance to assess the robustness of isotope signatures and thus consider the potential for clinical application of isotope-based cancer biomarkers. To address this issue, we took advantage of a combination of isotope, metabolome and lipidome analysis, and metabolism manipulation with DCA, to explore potential links between δ-values and metabolites. In practice, we used murine 4T1-Luc2 tumours representing triple negative breast cancer (TNBC) as weak DCA-responders, and V14 hormone receptor negative/HER2 positive tumours representing HER2 positive breast cancer (HER2+) as very good DCA-responders [61, 62]. Both tumour types are of BALB/c background (allowing growth in immune-competent hosts), and are p53-deficient (4T1 due to a frame shift mutation, and V14 due to loss of heterozygosity on a *Trp53^+/-^* background, i.e. p53 null) [63, 64].

We analysed tumour tissue and non-cancerous tissue in mice in which mammary tumours were induced via the injection of either of these cancer cell lines and looked at potential correlations between metabolism and isotope abundance, using multivariate analyses adapted to multi-omics datasets. Taken as a whole, our data show (*i*) the somewhat low δ^15^N values in tumours regardless of the tumour model, with a concurrent change in urea cycle indices (fumarate-to-aspartate ratio, urea-to-arginine ratio), and (*ii*) the generally high δ^13^C values linked to a lower content in triglycerides, regardless of the tumour model. DCA was effective in inhibiting tumour growth but did not impact on δ^13^C, while its effect on δ^15^N was dependent on the tumour model, reflecting important differences in background metabolism between different tumours, and the influence of N-containing lipids.

## Methods

### General mouse conditions

Animal experiments were conducted with the approval of the Australian National University Animal Ethics Experimentation Committee (Protocols A2011-008 and A2014_19) under the guidelines established by the Australian National Health and Medical Research Committee. Female BALB/c mice, were purchased from the ANU Animal Facility and were housed in small micro-isolator cages in a temperature-controlled environment with a 12 hr light / dark cycle. Autoclaved regular breeders mouse chow (Gordon’s Specialty Feeds, Bargo, NSW) was supplied ad libitum. BALB/c-*Trp53^+/-^* mice were bred and housed in the same facility.

### Cell lines for transplantation for mammary tumours

4T1-Luc2 (ATCC CRL-2539-LUC2) mouse mammary tumour cells were purchased from ATCC (VA, USA) and were grown at 37°C, 5% CO_2_ in RPMI-1640 media supplemented with 10% fetal bovine serum (FBS), 10 mM HEPES and 2 g/L NaHCO_3_. The V14 mammary tumour cell line was derived by Dr. Anneke Blackburn from a spontaneous mammary adenocarcinoma arising in a BALB/c-*Trp53^+/-^* mouse [64]. The original tumour and transplanted outgrowths from this tumour have previously been determined to be estrogen receptor and progesterone receptor negative, HER2/neu positive, and to have lost the wild-type allele of *Trp53*. This tumour line was not metastatic [61]. V14 cells were grown at 37°C, 5% CO_2_ in DMEM/F-12 medium supplemented with 25 mM HEPES (MP Biomedicals, USA), 1.2 g/L NaHCO_3_, 2% adult bovine serum (ABS), 0.1% PSN (3% penicillin, 5% streptomycin and 5% neomycin), 5 ng/ml epidermal growth factor (EGF), 10 μg/mL insulin, 15 μg/mL gentamicin (Sigma, MO, USA).

### In vivo tumour models

Immunocompetent female BALB/c mice, 8-14 weeks old (age-matched within each experiment), were anaesthetized with isofluorane and injected with prepared tumour cells in 15 µL of serum-free media into both left and right 4^th^ mammary glands. Mice bearing 4T1 tumours were treated with DCA at 50 mg/kg/day in a single i.p. injection. At the end of the experiment, mice were sacrificed by asphyxiation with carbon dioxide, and tumour tissue from the right side tumour collected for the analysis in this paper. Mammary gland tissue was also collected from these mice, pooled from 4^th^ (adjacent to tumour) and 5^th^ mammary glands on both sides. For mice bearing V14 tumours, DCA was administered in the drinking water at 1.5 g/L, and water consumption was monitored by weighing the water bottles every 2-3 days. The presence of DCA did not alter water consumption significantly. This delivered no more than 190-220 mg DCA/kg/day (which includes water wastage). While this is potentially a large difference between the two models in the actual dose of DCA, the 4T1 experiment included a group on 100 mg/kg/day i.p. which did not show any greater effect on tumour response (data not shown). Tumour size was monitored every 2-3 days with electronic callipers. The experiments were terminated when the tumour burden was too high in the control groups, (47 vs 28 days of growth for V14 and 4T1s respectively due to different growth rates). After sacrifice, the fresh tumour tissue was trimmed of non-tumour tissue and cut into several pieces, each weighing approximately 50-100 mg, placed in a cryotubes and snap frozen in liquid nitrogen for metabolic and mRNA analyses.

### Non-cancer tissues from Trp53^+/-^ mice and 4T1 tumour-bearing mice

Non-cancerous mammary gland tissue was collected from the 4T1 tumour-bearing mice described above (pooled from 4^th^ (adjacent to tumour) and 5^th^ mammary glands on both sides). Since the tumours were quite large, the mammary glands were quite depleted of fat compared to tumour-free mammary glands. Healthy mammary gland tissues (4^th^ mammary gland, with the lymph node excised) and non-mammary gland adipose tissues were also collected from 12 week old BALB/c-*Trp53^+/-^* female mice housed under the same conditions

### Isotope analyses by elemental-analysis/isotope ratio mass spectrometry (EA-IRMS)

Fresh tissue samples were frozen-dried (lyophilised). About 0.7 mg (dry weight) was weighed with a micro-balance (Ohaus Discovery DV215CD, Pine Brook, New Jersey, USA) into tin capsules (solids “light”, 5 × 9 mm, ThermoFisher Scientific, France). EA-IRMS analyses were carried out using a Flash EA elemental analyser coupled to a Delta V spectrometer run in continuous flow as previously described [54]. ^13^C and ^15^N natural abundance was expressed using the δ notation (δ^13^C and δ^15^N, in ‰ or mUr) as δ = (*R*/*R*_st_ – 1)·10^3^ where *R* is the heavy-to-light isotope ratio and *R*_st_ stands for the isotope ratio in the international reference (V-PDB for ^13^C and air N_2_ for ^15^N). The specific isotope composition of the net nutritional uptake (food ingested minus losses via excretion), could not, quite obviously, be known directly. In addition, food ingestion by mice varies, depending on their day-to-day preference and feeding behaviour. We therefore expressed δ-values relative to the background, time-integrated nutritional signature provided by liver. To do so, we used the isotopic difference Δδ (in ‰) = δ_sample_ – δ_liver_. For further explanations related to nutrition-based isotopic corrections, see [43].

### GC-MS metabolomics

Metabolomics were conducted by gas chromatography/mass spectrometry (GC-MS) on the Joint Mass Spectrometry Facility at ANU. Samples (5-10 mg dry tissue) were ground in liquid nitrogen and then a mixture made of 400 μL methanol, 400 μL chloroform and 360 μL H_2_O was added. Ribitol (7.5 nmol, dissolved in methanol) was added as an internal standard. After homogenisation and centrifugation at 15,000 rpm for 15 min at 4 °C, the aqueous phase was collected and centrifuged again. Aqueous phases were pooled, spin-dried under vacuum and stored at −80 °C until further analysis. GC-MS analyses were carried out as [65, 66] using derivatised extracts (trimethylsilylation) with methoxyamine and MSTFA (*N*-trimethylsilyl-*N*- methyl trifluoroacetamide) in pyridine, with the addition of an alkane mix to compute retention indices. Data were extracted with Metabolome Express [67] and normalised using the signal of ribitol and the dry mass of the sample.

### LC-HRMS lipidomics

Lipidomic analysis was performed on an arbitrary set of tissue samples that were homogenized in water at 100 mg/mL. The lipids were then extracted with methyl-*ter*-butyl ether (MTBE, Biosolve, Netherlands) as described previously [68]. Homogenates (50 μL) were successively mixed with 450 μL of ice-cold MTBE, 1500 μL of ice-cold methanol (Biosolve, Netherlands), and 375 μL of water. The mixes were centrifuged 10 minutes at 10,000 g at 4°C, and 800 μL of the supernatant was sampled. The samples were then dried under a nitrogen stream. Samples were finally resuspended in 150 μL acetonitrile/isopropanol/water (65/30/5, v/v/v) for liquid chromatography–high-resolution mass spectrometry (LC-HRMS). Quality control samples (QC) were prepared (n = 12) by pooling 20 μL of each homogenate. Samples and QC were arbitrarily randomized before analysis [69, 70] by LC-HRMS. LC-HRMS analyses were performed on a Synapt^TM^ G2 HRMS Q-TOF mass spectrometer equipped with an electrospray ionization (ESI) interface operating in the positive and negative modes and an Acquity H- Class^®^ UPLC^TM^ device (Waters Corporation, Milford, MA, USA). Lipid separation was achieved after injection of samples (10 µL) onto an Acquity^®^ CSH C_18_ column (2.1 mm × 100 mm, 1.7 µm; Waters Corporation) held at 55°C. The mobile phase was composed of an acetonitrile/water (60/40, v/v) mixture as solvent A and an isopropanol/acetonitrile (90/10, v/v) mixture as solvent B, each containing 10 mM ammonium acetate and 0.1% formic acid (Biosolve). The elution was carried out using a multistep gradient of solvent B in solvent A over 22 min at a constant flow rate of 400 µL/min as described in [71]. The HRMS mode was applied for lipid detection (mass-to-charge ratio (*m/z*) range 50-1,200) at a mass resolution of 25,000 full-widths at half maximum. The following parameters were used for ionization process: capillary voltage, +2 kV (for both positive and negative ionization modes); cone voltage, 30 V; desolvation gas (N_2_) flow rate, 900 L/h; desolvation temperature, 550 °C; and source temperature, 120 °C. Leucine enkephalin solution (2 µg/mL in 50% acetonitrile containing 0.1% formic acid) was infused at a constant flow rate of 10 µL/min in the lockspray channel, allowing for correction of the measured *m/z*. Mass correction was applied throughout the batch at *m/z* 556.2771 (ESI+) and 554.2616 (ESI-). Data acquisition and processing, including peak detection, integration, alignment, and normalization, were achieved using MassLynx^®^ and MakerLynx^®^ software (version 4.1, Waters Corporation). Lipid markers were extracted from the detected variables with an in-house database built by the use of lipid standards, their exact mass measured (taking into account the specifically formed adduct ions), their elemental compositions with a mass error below ± 5 ppm, their retention times and their fragmentation patterns as described in many studies [72–74]. Simultaneously, QC sample analyses were executed to evaluate the analytical system performance during the analytical run. The relative standard deviation (RSD, %) was calculated for peak areas to highlight the repeatability of the analytical process. Finally, selected lipid markers having a RSD value below 30% were retained for multivariate analysis [69].

### Statistical analyses

Integration of the combined isotope, metabolome and lipidome dataset was carried out using consensus orthogonal partial least squares (C-OPLS), which is a supervised multiblock method allowing the combined statistical analysis of multiple data sources using a single multivariate model [75]. Monte Carlo Uninformative Variable Elimination-Partial Least Squares (MCUVE- PLS) was performed based on an ensemble of 10^4^ models and a ratio of training samples of 0.7 using the libPLS 1.98 toolbox [76]. A threshold of reliability index value of 1.00 was applied to remove variables considered as uninformative. C-OPLS and MCUVE-PLS analyses were performed under the MATLAB® R2022a environment (MathWorks, Natick, USA), while corresponding figures were generated with R (version 4.1.0) and RStudio (version 2023.03.0, +386). Simple linear regression (univariate analysis) was computed using OriginPro2020. Adjusted *P*-values were computed using the Benjamini-Hochberg false discovery rate procedure [77]. (Further details of each analysis can be found in Supplementary Material.) All metabolomics and lipidomics data were normalized to the specific mass of tissues and some variables such as ratios and sums were calculated and integrated in the data set. (The dataset used and/or analyzed for this study is available upon reasonable request.)

### Nanostring mRNA analysis

At the time of sacrifice, pieces of tumour tissue each weighing approximately 50-100 mg were snap frozen in liquid nitrogen for future metabolic and mRNA analyses. Tissue was dissociated using the GentleMACS Octo and M tubes (Miltenyi Biotec) and RNA was extracted using the Maxwell® RSC simplyRNA Tissue Kit (Promega, Madison, USA) according to the manufacturer’s instructions. Gene expression analysis was undertaken using the NanoString nCounter analysis system, (Bruker Spatial Biology, Seattle, WA) using the commercially available Mouse Metabolic Pathways Panel and the Mouse PanCancer Pathways Panel. Samples were hybridised with the panel probes for 20 hours, processed on the NanoString Prep Station and the target probe complex immobilised onto the analysis cartridge. The cartridges were scanned by the nCounter Digital Analyser at 555 fields of view.

Gene expression data was analysed using the Advanced Analysis Module in the nSolver™ Analysis Software (version 4; NanoString Technologies, WA, USA) and R packages including NanoStringNCTools (3.19) and limma. The advanced analysis module enables quality control assessment and thresholding. NanoStringNCTools was used to assess various sample integrity against technical QC metrics, including lane binding density, lane fields of view, external RNA controls consortium linearity (ERCC) and ERCC-LOD. Samples with a low percentage of housekeeping genes or binding density exceeding 2.2 were flagged for inspection in the downstream analysis. Housekeeping probes to be used for normalisation were selected using the geNorm algorithm in the NormqPCR R library while technical variation was normalised through internal controls. Cluster analysis, differential gene expression (DGE), Pathview Plots and immune cell profiling between the groups was undertaken with NanoStringNCTools. P-value adjustment was performed using the Benjamini-Hochberg method of estimating false discovery rates (FDR).

## Results

### Isotope composition in tumour and non-tumour tissues

The % content of N or C varies with tissue composition – protein and nucleic acid rich tissues such as liver will have high N content, while lipid rich tissues such as adipose and mammary gland will have high C content. Liver and tumours are typically protein-rich and lipid-poor tissues, and are therefore expected to have higher %N and lower %C compared to adipose and mammary gland tissues. In general, this was what we observed in our sample groups. The %N and %C in tumours (Figure 1A and B) were similar to those measured in mouse liver (10.8±1.3 %N, 48.5±4.3 %C, mean ± SD of all 28 mice combined). Compared to the lipid-rich adipose and mammary glands, tumours contained more N (Figure S1A) and were, on average, less rich in C (Figure S1B). Interestingly, %N was significantly different between the two tumour models (Figure 1A), while %C was not (Figure 1B). Tumours were enriched in nitrogenous compounds, as shown by the significantly higher %N (Figure 1A), with V14 tumours having the highest %N. This effect was due to either lower cellular N turnover rate, higher content in proteins or N-containing metabolites, or lower stored lipids content (which are mostly devoid of N). Conversely, tumours were on average less rich in carbon (Figure 1C), likely reflecting the lower amount of lipids (and thus adipose tissue) which are C-rich compounds (70 to 80% carbon).

**Figure 1.**
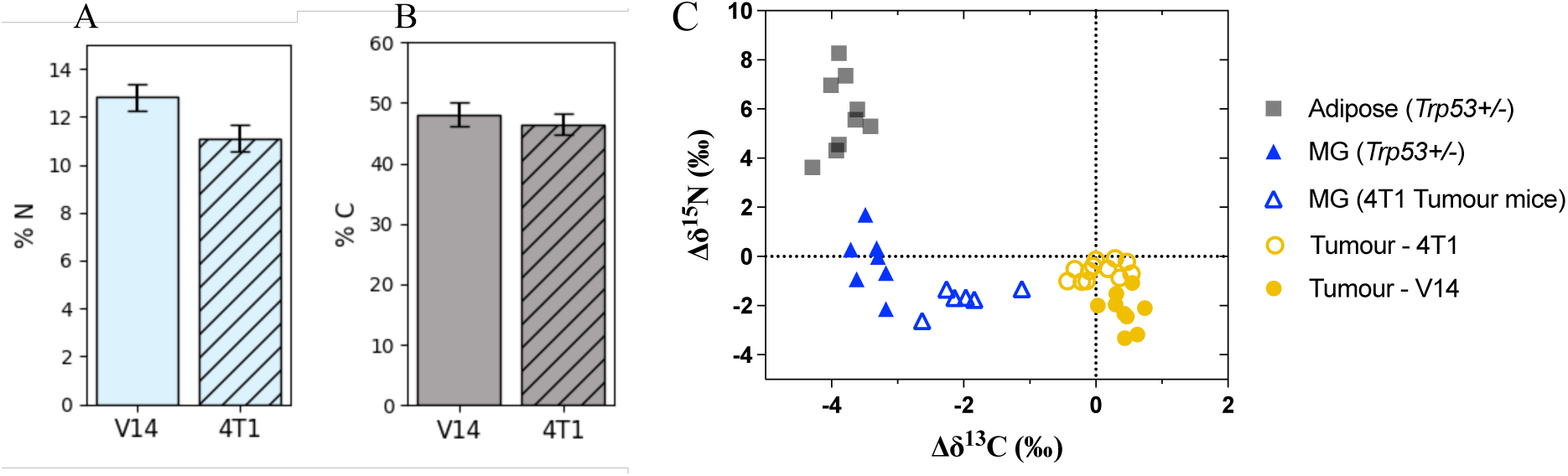
N and C content and isotope composition in mammary tumour, adipose and mammary gland (MG) tissues. **(A)** Nitrogen content (% of dry weight) is different between the V14 and 4T1 tumours (*P* < 0.01), but **(B)** carbon content (% of dry weight) is not. **(C)** Isotope biplot showing the natural abundance in ^13^C (Δδ^13^C) and ^15^N (Δδ^15^N) which can distinguish between adipose, mammary gland and tumour tissues.

The natural abundances of the isotopes ^13^C (Δδ^13^C) and ^15^N (Δδ^15^N) (liver background-subtracted) are shown in Figure 1C. The Δδ^13^C and Δδ^15^N values could discriminate between tumour and normal samples (mammary gland or adipose tissue). Tumour samples were significantly ^13^C-enriched by up to 4‰ compared to adipose tissue or mammary gland from *Trp53+/-* mice, regardless of tumour model, and by about 2‰ compared to mammary tissue adjacent to 4T1 tumours. Mammary glands and tumours tended to be relatively ^15^N-depleted (lower Δδ^15^N) compared to adipose tissue (Figure 1C). The Δδ^15^N value in V14 tumours was found to be lower than in 4T1 tumours (Figure 1C), with V14 tumours being significantly ^15^N- depleted compared to any other tissue or tumour (*P*<0.05).

### Multivariate metabolic signature of tumour vs non-tumour tissues

A combined (multiblock) C-OPLS discriminant analysis (DA) of the three datasets (215 features in total: 7 isotopic features, 96 metabolomic features and 112 lipidomic features) was conducted with samples grouped for comparison of tumours (*n* = 21) with non-tumour samples (*n* = 23), regardless of the tumour type, mouse treatment or mouse genotype (validation details in supplementary material – Figure S2A). A clear separation between tumour and non-tumour samples was observed along the predictive component (x-axis, Figure S3A), explaining 31% of total variance. The block contributions (Figure S3B) underlined the major importance of isotopic features (41%), followed by metabolomic (31%) and lipidomic features (28%) to the predictive component. The orthogonal component (y-axis, Figure S3A) differentiated between the two different tumour models, and also between the three different non-tumour sample groups. The orthogonal component explained 43% of total variance, spread over metabolomic (39%), lipidomic (44%), and isotopic features (17%) (Figure S3C). Individual feature contributions were investigated using the combined loading plot (Figure S3D). In agreement with Figure 1C, tumours were associated with a lower Δδ^15^N (dark green frame) and higher Δδ^13^C value (dark blue frame) compared to the non-tumour samples. These results confirm that in our context, isotope composition can distinguish between mammary normal and tumour tissues in mice.

In addition to isotopic differences, tumours were enriched in many metabolites such as succinate, some amino acids and their derivatives (phenylalanine and hydroxyproline), and sugar derivatives (ononitol and threonate), likely reflecting typical metabolic changes that occur in cancer cells, such as increased glycolysis (the Warburg effect) and increased glutamine utilisation (glutaminolysis) [1].

There were also differences in lipid composition between the normal and tumour tissues, with a general increase in many molecular species of phosphatidyl choline (PC, membrane lipids), changes in cerebrosides (CB) and sphingomyelins (SM) (signalling lipids) and depletion in oleate (18:1 fatty acid). A decrease in triglycerides (TG) was also evident, consistent with the generally lower lipid content of tumours compared to adipose and mammary gland tissues, since TGs are the major form of lipid accumulation in these tissues. Tumours were also enriched in arginine and ornithine, enriched in C_4_ acids, and had a low fumarate-to-aspartate (Fum/Asp) ratio, consistent with modifications to the urea cycle and mitochondrial activity. (Note that Σ_C4 metabolites includes succinate, fumarate, malate and aspartate. Oxaloacetate could not be measured in our analysis.)

### Potential metabolic drivers of isotope composition

Potential links between metabolic features and isotope abundance were explored via a covariation analysis between Δδ values and metabolomic and lipidomic feature peak areas across all tumour and non-tumour tissues. (Validation details in supplementary material – Figure S2B and S2C.) Volcano plots were used to identify the best potential drivers of Δδ^15^N and Δδ^13^C (Figure 2). Δδ^15^N correlated most strongly with cerebroside-to-sphingomyelin ratio (CB/SM) and Fum/Asp (-logP >4.5), sum of triglycerides/total lipids (Σ_TG/tot), sum of dihexosyl CBs/total lipids (Σ_dihexCB/tot), urea-to-arginine ratio (Urea/Arg), and oleate (- logP >2.5). Δδ^15^N was inversely correlated to phosphatidyl and lysophosphatidyl choline (PC and lysoPC) molecular species (Figure 2A). Δδ^13^C was positively correlated to PC and lysoPC species, succinate (Succ), fumarate (Fum) and Σ_C4 (-logP >7.5), and inversely correlated to TGs (-logP >14), CB/SM, Fum/Asp and oleate (-logP >5) (Figure 2B).

**Figure 2.**
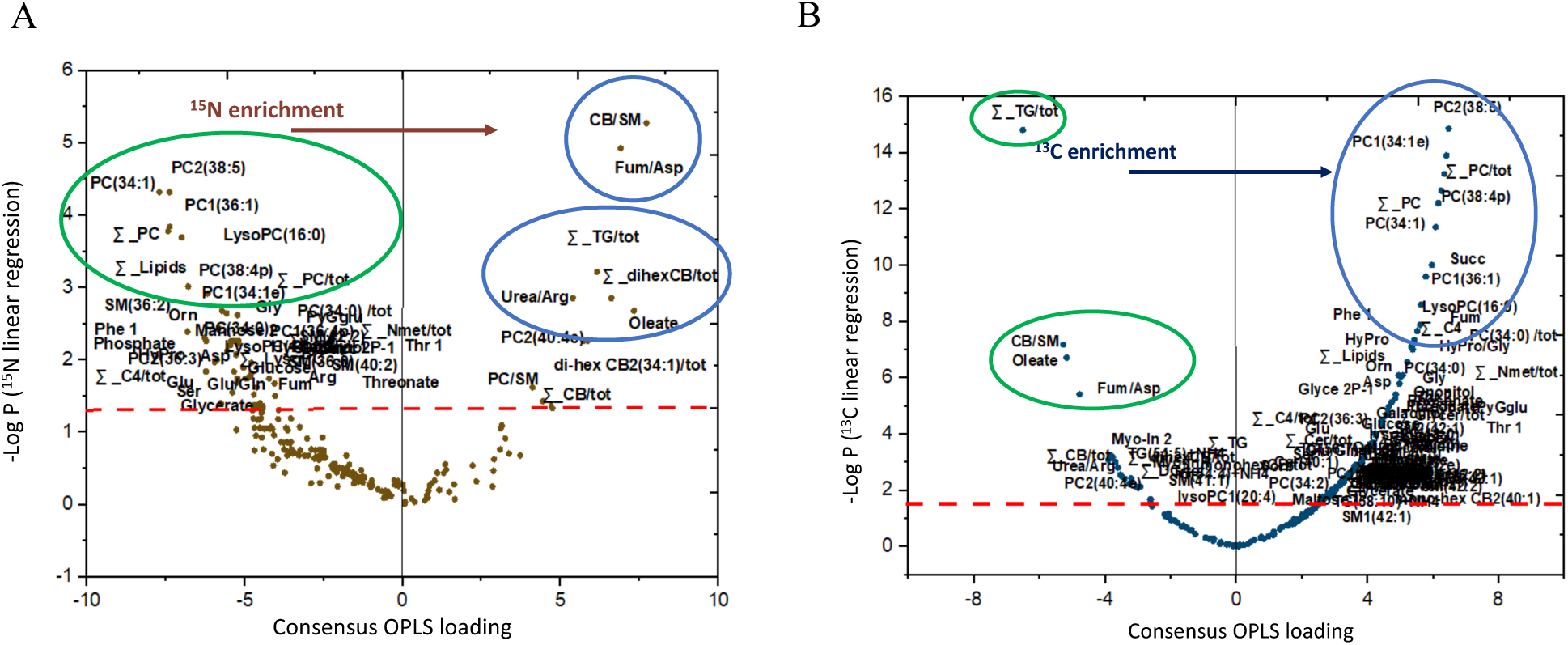
Potential drivers of isotope composition: metabolite and lipid features that correlate with natural isotope abundance in tumour and normal tissues combined. The volcano plots identify the metabolite or lipid features which most significantly correlated to the **(A)** Δδ^15^N and **(B)** Δδ^13^C across all tumour and non-tumour tissues (n=21 and 23 respectively, 44 tissues in total). Volcano plots were constructed combining C-OPLS loadings (p_corr_) with *P*-values computed from linear regression (after Benjamini-Hochberg false discovery rate correction) between features and Δδ values. The red dashed line represents *P=*0.05. Features with *–*log(*P*) above 1.33 (*P*<0.05) were annotated. Positive correlations (blue ellipses), inverse correlations (green ellipses).

### Effect of tumour model and DCA on natural isotope abundance in tumours

The two mouse mammary tumour models under investigation represent two human breast cancer subtypes: 4T1 is triple negative (TNBC) and V14 is HER2 positive (HER2+). They differ in their growth rates and their response to the metabolic modulator, DCA (Figure 3A). The average tumour size of rapidly growing 4T1 control and DCA-treated tumours was significantly different from day 16 onwards, however, after a small delay they show almost no change in growth rate in response to DCA (tumour doubling time of 6.9 and 7.2 days, control and DCA-treated respectively, Figure 3A inset). In contrast, slower growing V14 tumours show a robust growth inhibition when treated with DCA, with the tumour doubling time slowing from 10.9 to 19.6 days.

**Figure 3.**
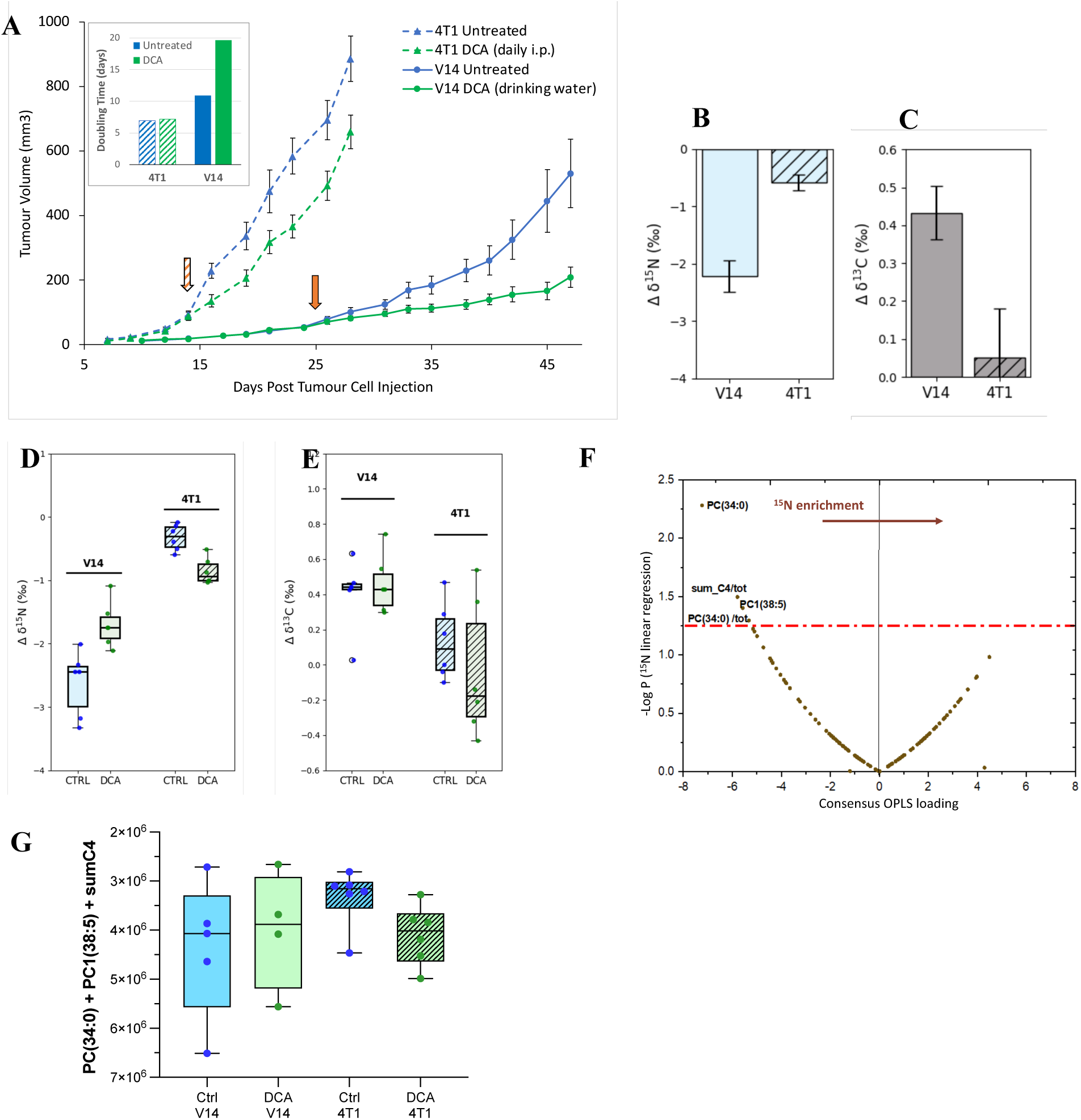
Isotopomic differences and drivers in mammary tumours, untreated and treated with DCA. **(A)** Tumour growth curves for 4T1 and V14 mammary tumours, untreated and treated with DCA (mean ± SEM, n=8-16 tumours per group). Orange arrows indicate start of DCA treatment. Inset shows the tumour doubling time according to an exponential curve fit from day 14 (4T1) or day 24 (V14) to the end of the experiment. **(B) and (C)** The natural abundance of ^15^N (Δδ^15^N) is depleted and of ^13^C (Δδ^13^C) is enriched in both tumours, but to a different extent (mean ± SEM, n=4-6 tumours per group. *P* < 0.0001 for Δδ^15^N and *P* < 0.005 for Δδ^13^C). **(D)** DCA treatment significantly changed the Δδ^15^N of both tumour models (*P* = 0.024 for V14 and *P* = 0.001 for 4T1) (p<0.05), but in opposite directions, and to different extents. **(E)** DCA treatment did not significantly change the Δδ^13^C of either tumour model. (Median, quartiles, and range displayed. n=4-6 tumours per group). **(F)** Drivers of Δδ^15^N values across all 4 tumour groups combined were identified with a volcano plot. The red dotted-dashed line represents the threshold of *P=*0.05. **(G)** Sum of the metabolomic / lipidomic values of the Δδ^15^N drivers identified in (F).

The isotope biplot of Δδ^15^N and Δδ^13^C (Figure 1C) could distinguish between the two tumour models. Examining this more closely, we found that both tumour types were Δδ^15^N depleted and Δδ^13^C enriched, but to different extents (Figure 3B and 3C). To examine this further, the isotope composition of the four tumour and treatment groups was analysed (Figure 3D and 3E). Surprisingly, there was an opposite change in Δδ^15^N in response to DCA treatment in the two tumour models (Figure 3D). DCA increased Δδ^15^N significantly by ∼0.9‰ in V14 tumours, whereas 4T1 tumours showed a significant decrease in Δδ^15^N of ∼0.5‰. In contrast, there was no significant effect of DCA on Δδ^13^C in either tumour model (Figure 3E). To identify metabolic features that contribute to the cellular content of ^15^N in tumours, C-OPLS regression analysis of a 50-feature multiblock dataset was undertaken on tumour samples only with Δδ^15^N as the Y quantitative response variable (analysis and validation details in supplementary material). The resulting volcano plot indicated that across all the tumour samples, only 4 features were robustly associated with Δδ^15^N, with inverse correlations – PC(34:0) (alone, or as a percentage of total lipids), PC1(38:5), and Σ_C4/tot (Figure 3F). These features are consistent with those found across tumour and normal tissues combined (Figure 2A), although the small number of features identified contrasts starkly. To confirm these metabolites could account for the changes in Δδ^15^N, the sum [PC(34:0) + PC1(38:5) + Σ_C4] for each tumour was calculated (Figure 3G). The resulting pattern of differences between the tumour groups paralleled the Δδ^15^N values, supporting that changes in the levels of these features could be driving the differences in Δδ^15^N values. Thus, Δδ^15^N values were able to point to underlying metabolic differences that associated with differences in tumour growth responses to DCA *in vivo*.

### Metabolic alterations associated with DCA treatment or cancer subtype

To explore the effect of DCA and tumour model on metabolism more broadly, a multivariate analysis was carried out. To improve the performance of the C-OPLS-DA model, mid-level data integration analysis and uninformative variable elimination (UVE) were done (validation details in supplementary material, Figure S2D). The resulting (trimmed) combined dataset (comprised of 1 isotopic feature, 47 metabolic features and 57 lipid features, total 105) was then subject to C-OPLS-DA analysis. The score plot indicates the four tumour groups were clearly separated by this analysis (Figure 4A). The overall effect on tumour metabolism of the tumour model was much stronger than the effect of DCA treatment (explaining 41% and 10% of total variance, respectively). DCA was associated with the predictive component while tumour models were associated with the orthogonal component (Figure 4A). Block contributions showed the importance of metabolites (43%), followed by lipids (40%) and then isotopes (17%) in discriminating between untreated and DCA-treated tumours (Figure 4B). Similar block contributions were observed for discrimination between 4T1 and V14 tumours (Figure 4C).

**Figure 4.**
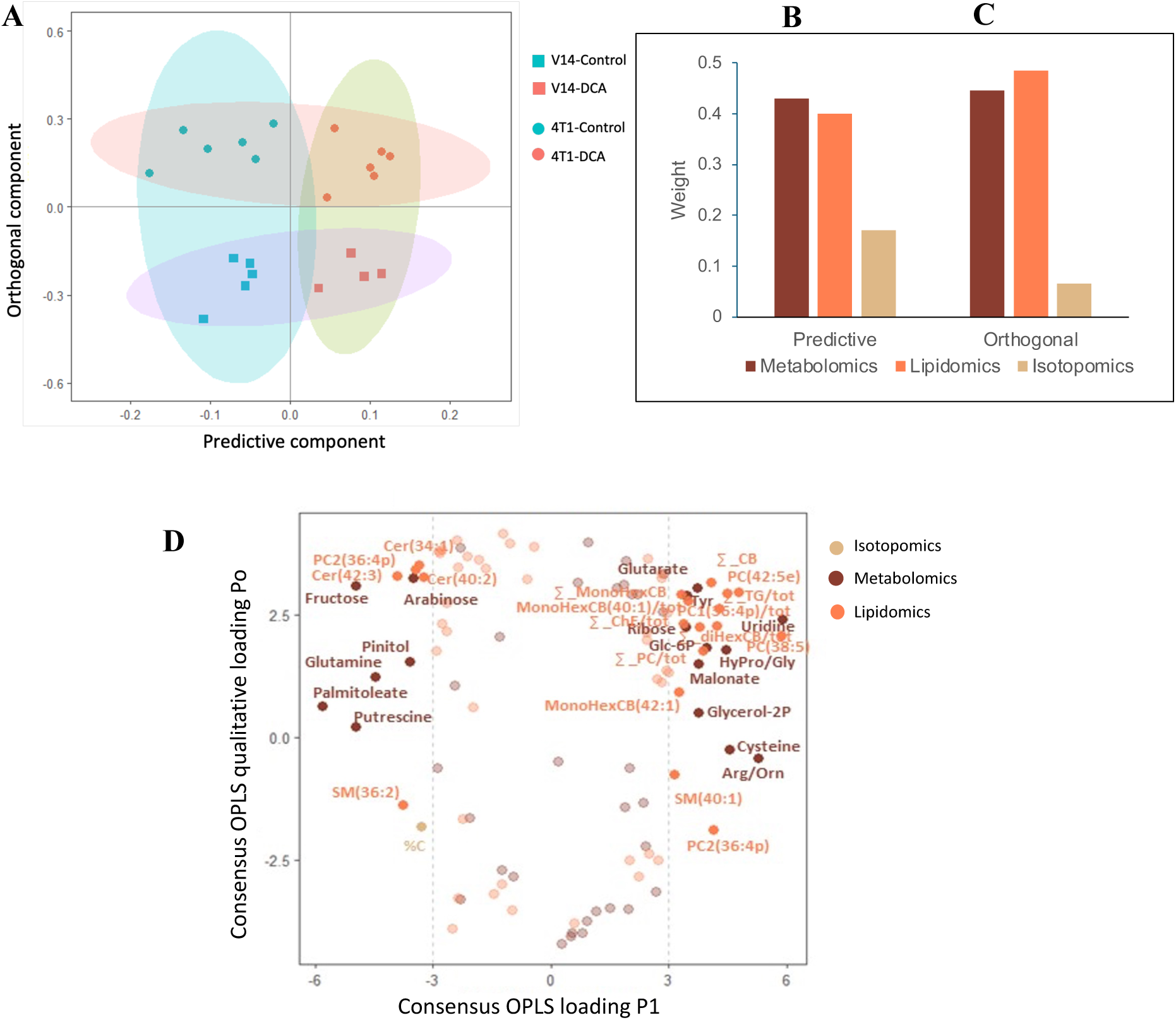
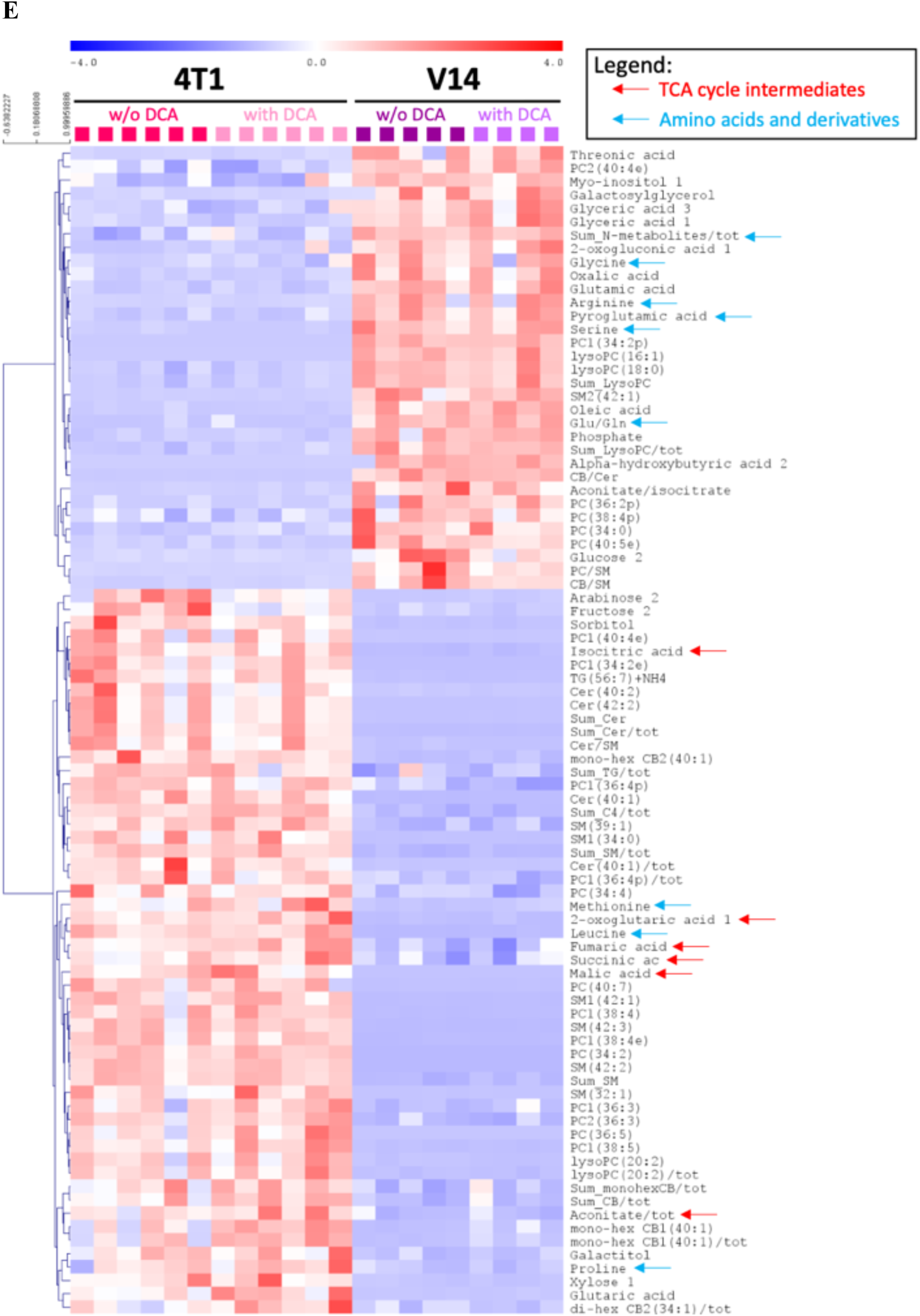
Differences in combined metabolomics, lipidomics, and isotopic patterns in tumours, untreated and treated with DCA. **(A)** Score plot of the C-OPLS-DA (R²Y = 0.83 and Q²Y = 0.41) discriminates the DCA-treated tumours from the control tumours (predictive component) and also between V14 and 4T1 tumour models (orthogonal component). The ellipses are for a normal distribution with a level of 0.9. **(B)** and **(C)** Block contributions showing the importance of metabolites, lipids and isotopes in sample discrimination **(B)** with vs. without DCA treatment, and **(C)** tumour type discrimination. **(D)** The C-OPLS loading plot shows the significant features among isotopomics, metabolomics and lipidomics data that discriminate DCA-treated from untreated tumours. The variables are color-coded according to which block they belong: isotopomic, lipidomic or metabolomic block. **(E)** Most significantly different features (combined metabolomics and lipidomics) between tumour models (two-way ANOVA). Features with P < Bonferroni threshold are shown. Metabolites mentioned in text are labelled with coloured arrows.

The impact of DCA on metabolism is shown in the C-OPLS loading plot (Figure 4D). This shows that DCA caused changes in a variety of tumour metabolites, beyond a simple reversal of the glycolytic phenotype. The significant changes (n=36 altered features across multiple blocks) included an increase in total triglycerides and cerebrosides (along with a complex effect on phospholipids), some amino acids (tyrosine, hydroxyproline, cysteine), nucleotide precursors (uridine, ribose), and glucose 6-phosphate. There were also decreases in sugars/sugar alcohols (fructose, arabinose, pinitol), and a change in polyamine/urea cycle metabolites, with an increase in the arginine-to-ornithine ratio and a decline in putrescine content.

In contrast, the number of significantly different metabolic features between the two tumour types was much greater (n= 87) (Figure 4E) than those altered by DCA. This is consistent with the relative contributions to total variance calculated from the C-OPLS-DA analysis. 4T1 tumours were enriched in TCA cycle intermediates (such as isocitrate, 2-oxoglutarate, fumarate and malate), leucine and methionine, ceramides and sphingomyelins while V14 tumours were enriched in total lysoPCs and nitrogenous compounds such as serine, pyroglutamate and glutamate, with a higher glutamate-to-glutamine ratio. This highlights how the metabolic difference between the two tumour types is much greater than the differences due to DCA treatment.

### Interaction effects between DCA treatment and tumour model

An additional analysis was performed on the combined metabolomics and lipidomics data looking for an interaction effect – that is, an interaction between tumour model (V14 or 4T1) and the response (or lack of response) to DCA treatment. Six features were identified (Figure 5). Most strikingly, 4 of these 6 features show dramatically different levels between the tumour models, with malate, fructose, malonic acid and glutaric acid being almost totally absent in V14 tumours compared to 4T1 tumours. 4T1 tumours were able to induce or maintain higher levels of fumarate, malate, malonic acid, glutaric acid and di-hex CB2(34:1) in response to DCA, where V14 tumours did not. Conversely, 4T1 palmitelaidic acid levels were decreased by DCA, whereas V14 tumour levels were unchanged.

**Figure 5.**
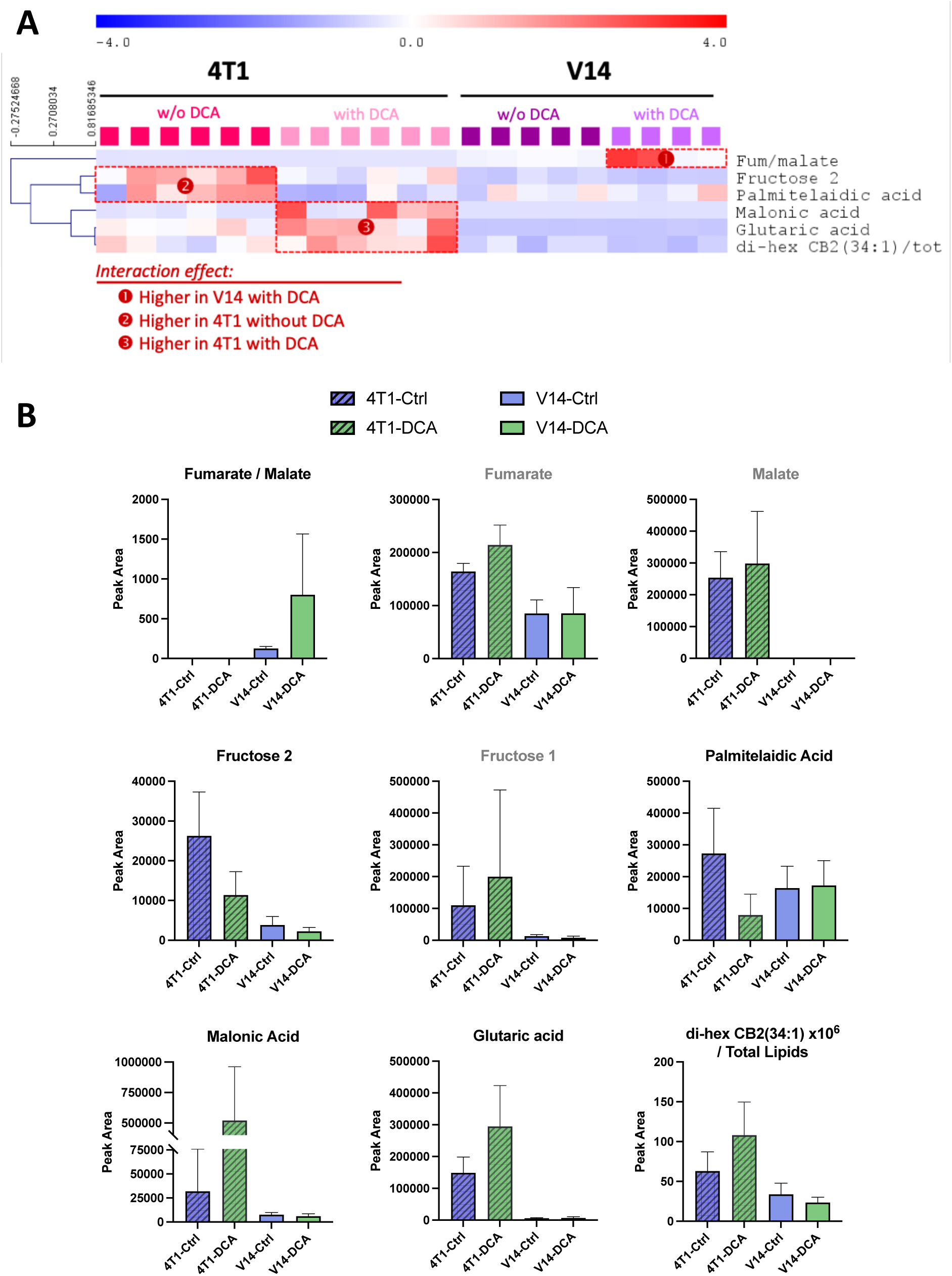
Interaction effects between tumour model and DCA treatment. **(A)** Features that respond differently to DCA treatment between 4T1 and V14 tumours (two-way ANOVA, *P*<0.05). **(B)** Mean +/- SD for features displayed in (A). Grey headings display closely related features that did not reach significance.

### Gene expression changes after DCA treatment correlate with metabolite changes

To complement the metabolic analysis, the transcriptome of the DCA-sensitive V14 tumour was analysed on the Nanostring Metabolic Pathways panel for gene expression changes due to DCA treatment. Of the 702 genes that were examined, no genes showed differential expression (DE) using the standard statistical criteria. However, over several weeks of treatment with DCA, even subtle changes in mRNA expression may have significant biological effects, so we examined the data with lower stringency, considering unadjusted p values < 0.1. Only 29 genes (Table S2 and S3) showed DE between the untreated and DCA-treated V14 tumours (p < 0.1), despite this treatment being capable of dramatically slowing tumour growth (Figure 3). The 29 genes were examined for GO pathway enrichment (https://www.pantherdb.org/). Only one major GO pathway / hierarchy was identified, that of Carboxylic acid metabolic process (Table S2). The 29 genes were also manually sorted into 4 categories based on the gene annotations: metabolic / enzymatic activity (n=13), binding proteins (n=7), transcription factors (n=3) and immune-related (n=6) (Figure 6, Table S2 and S3). Increases in expression ranged from 12 to 223% while decreases ranged from 12 to 54% (Figure 6A), changes likely to be of biological significance over weeks of DCA treatment. Consistent with the interaction effect features, the DE genes were involved in diverse aspects of metabolism, reflecting the centrality of the PDH/PDK axis in influencing metabolism overall.

**Figure 6.**
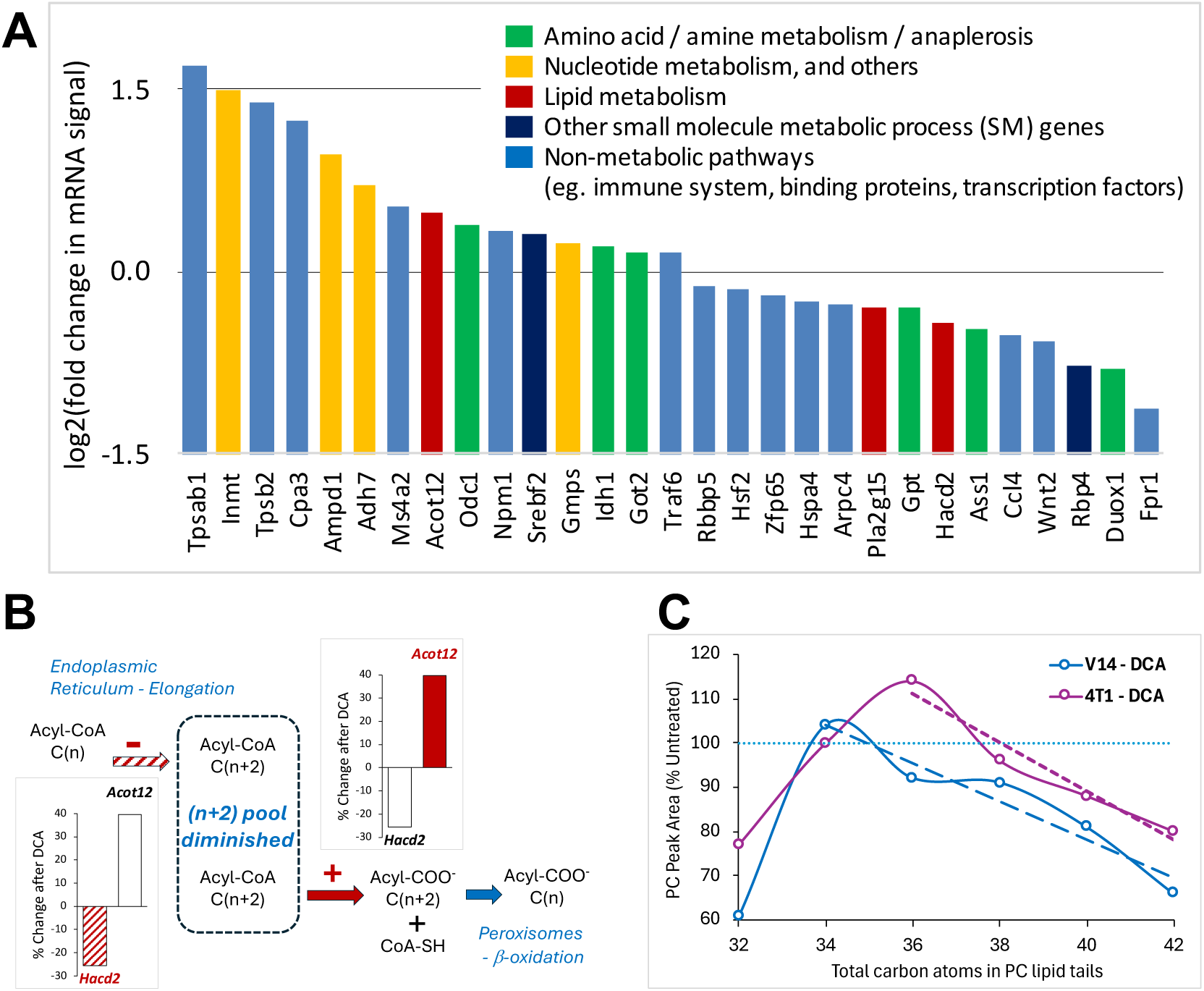
Gene expression changes after DCA treatment correlate with metabolomic features. **(A)** Differentially expressed (DE) genes in V14 tumours after DCA treatment (Nanostring Mouse Metabolism Panel). **(B)** Schematic showing *Hacd2* and *Acot12* (genes involved in fatty acid elongation and oxidation, respectively) and how the expression changes might be expected to alter the cellular fatty acid pool. **(C)** Linear regression of changes in PC content after DCA treatment, expressed as a % of untreated tumour PC content (100% = no change with DCA treatment, blue dotted line). PCs were grouped according to the total number of C atoms in the lipid tails. *P*=0.007 and 0.024 for V14-DCA and 4T1-DCA respectively, for the slope being significantly non-zero (simple linear regression, Graphpad Prism). SM = small molecule metabolic process. PC = phosphatidyl choline.

Evidence of biological impact from these gene expression changes was sought in the metabolic data, focusing on lipid metabolism (Figure 6B). *Hacd2,* a gene involved in fatty acid elongation in the endoplasmic reticulum, showed decreased expression while *Acot12,* involved in hydrolysis of acyl-CoAs into free fatty acids for peroxisomal β-oxidation, was increased. Significant changes in these activities would be expected to result in a diminished pool of longer chain fatty acids and their derivatives. We examined phosphatidyl cholines (PC), the most abundant and diverse lipid components detected in our untargeted lipidomics, for evidence of shortening of the lipid tails after DCA treatment. Comparing the abundance of PCs according to the number of carbon atoms in their tails, DCA treatment was found to significantly decrease the amount of longer tailed-PCs (Figure 6C) compared to untreated tumours (100% blue dotted line), consistent with the schematic in Figure 6B. The effect was evident in both tumour models but was more pronounced in V14 tumours (effect seen for 35Cs and above vs 38Cs and above, V14 and 4T1 respectively), correlating with the efficacy of DCA *in vivo*. This may also relate to the difference in malonic acid levels between the two models with and without DCA treatment (Figure 5B), an important precursor for fatty acid synthesis in both the cytosol and mitochondria.

## Discussion

Natural abundance isotopomics is an emerging analysis tool for discovery of biomarkers of diseases such as cancer. Here we demonstrated that the combination of Δδ^15^N and Δδ^13^C was able to discriminate (i) between five different mouse tissue types (isotope biplot Figure 1C), (ii) between two different mammary tumour models (Figure 3B and 3C), and also (iii) between the responses of these two different models to treatment with the metabolic modulator, DCA (Figure 3E and 3F). Our isotopic analyses support earlier work on human biopsies of breast cancer, where Δδ^15^N and Δδ^13^C were able to differentiate between adjacent non-cancer breast tissue and cancer tissue [54]. Human breast cancer biopsies were significantly ^13^C enriched, corresponding to our findings in mice, and tended to be ^15^N depleted [54]. In several other cancer types, reviewed in [43] and more recently, in bladder and endometrial cancer [78, 79], Δδ^15^N and Δδ^13^C were also able to discriminate between normal and cancer human biopsies. In the case of bladder cancer, ^13^C depletion in the normal urothelium of bladder cancer patients was associated with a worse prognosis [78]. Thus, this work overall supports continuing investigation of natural abundance isotopes as potential biomarkers in cancer.

It is important to ask what natural abundance isotopes can tell us about the underlying biology and biochemistry of the samples analysed. The previous reports on Δδ^15^N and Δδ^13^C in cancers have in some cases correlated the changes with pathological and patient features, such as histological grade and disease-free survival, but little work has been done on exploring the metabolic changes driving the changes in natural abundance of isotopes in cancer [43]. In this mouse study, we performed combined analyses of untargeted metabolomics and lipidomics with the isotopomics to gather evidence of the metabolic pathways underlying the isotope composition of normal and tumour tissues (Figure 2, Figure S3 and Figure 3). Examining the correlation between Δδ^15^N and Δδ^13^C and individual metabolite levels across all the tumour and non-tumour samples combined found that there were many species with significant correlations, positive or inverse (Figure 2), including many lipid species, particularly PCs and total TGs. This study used lipid-rich normal tissues as a comparison point for the tumours, and lipids are known to be ^13^C-depleted compared to total biomass [69], thus our ability to robustly detect common lipid species as drivers of isotope composition across the tumours and adipose-rich normal tissues is as expected. When considering only tumours, the two tumour models were significantly different in both Δδ^15^N and Δδ^13^C (3B and 3C), but while Δδ^15^N was significantly changed by DCA treatment in both models, Δδ^13^C was not changed in either model (3D and E). This δ^13^C result was remarkable, given our understanding that DCA alters glycolysis [21], and strongly suggests that the ^13^C content in these tumours is independent of the glycolytic flux, but rather depends on the content in end products such as lipids (PCs and TGs) or C_4_ acids (fumarate, succinate, malate and aspartate in this analysis).

We focused our attention on drivers of ^15^N content in tumours, which highlighted only a few features, namely the phospholipids PC(34:0), and PC1(38:5), and Σ_C4 metabolites (Figure 3F). N-containing lipids such as PCs are naturally ^15^N-depeleted compared to total cellular material [80], and phosphoethanolamine head groups of polar lipids have effectively been found to be ^15^N-depleted by up to 8‰ compared to cellular material [81], so it makes sense that these features underly the ^15^N content of tumour tissues. Indeed, in our data, considering those features, the pattern of differences in δ^15^N across the two breast cancer models with and without DCA treatment was approximately recapitulated when the metabolomic / lipidomic values of those features were combined (Figure 3G). Breast cancers are well-known to be enriched in phosphatidyl choline and associated metabolites, mostly due to a change in the activity of phospholipases and choline kinase [82–89]. Lipidomics studies conducted so far have shown an increase in specific sphingomyelin molecular species in breast cancer cells (reviewed in [90]). Breast cancer cells have also been shown to have high activity of the enzyme UDP-galactose:ceramide galactosyltransferase. This enzyme synthesises galactosyl- based cerebrosides from ceramide, depleting ceramide and thereby impeding ceramide-mediated apoptosis induced by doxorubicin [91] and favouring drug resistance [92]. Other correlating drivers of ^15^N content included specialised lipids (di-hexCB2(34:1), CB/SM), oleate (18:1), and the C4 acids Fum/Asp, and succinate which differed between V14 and 4T1 tumours. These species play roles in lipid signalling, structure and storage, anaplerosis and the urea cycle.

Increased glycolysis (the ‘Warburg effect’ or ‘glycolytic phenotype’, yielding lactate) and increased glutamine utilisation (‘glutaminolysis’, feeding the TCA cycle) represent two well-known metabolic hallmarks found in many cancers [1], including in TNBC or HER2+ BrCa [93]. Further, it is well established that different breast cancer subtypes display different metabolic phenotypes. Our two mouse models represent different subtypes of breast cancer: V14 tumours are ER-PR-HER2+ (HER2+), while 4T1 tumours are ER-PR-HER2- (TNBC). TNBC typically display higher glycolytic activity and low mitochondrial respiration [93, 94], while both TNBC and HER2+ BrCas display elevated glutaminolysis [95], with the latter tending to have higher glutamine and lipid metabolism [93–96]). Such metabolic modifications are believed to be mediated by oncogenes (e.g., *c - Myc*) and/or tumour suppressors (e.g., *p53*) [1]. Both of these mammary tumour models do not have wild-type p53 (V14 is null, 4T1 is mutant), whereas *c - Myc* is amplified and overexpressed in 4T1 cells but its mutation status / expression level in V14 cells is unknown [61, 63, 64]. The models will vary in the profile of other mutations that influence their metabolism and biology, thus it remains to be seen how generalizable the isotope and DCA response results are for human breast cancer subtypes.

Since DCA tends to revert the glycolytic phenotype, it was anticipated that, alongside the effect on tumour growth (Figure 3A), DCA treatment would cause a general decrease in glycolytic intermediates availability (including pyruvate), impacting on several biosynthetic pathways [28, 29, 40]. Our data show a rather modest effect of DCA on metabolic features, explaining about 10% variance only (Figure 4A). Various pathways appeared to be affected, such as lipid synthesis (with, e.g., an increase in glycerol 2-phosphate and malonate and a decrease in palmitoleic acid) and amino acid metabolism (Figure 4D). Also, DCA directly impacted on sugar metabolism, with a concurrent increase in glucose 6-phosphate and decrease in fructose, reflecting a change in the balance between glycolysis and the pentose phosphate pathway [30].

We have validated the biology of our metabolomic analysis by complementing it with findings from gene expression changes in V14 tumours. DCA treatment altered mRNA expression of two genes involved in lipid metabolism, increasing *Acot12* and decreasing *Hacd2* expression, which was associated with shorter lipid chains in the PCs of both tumour models (Figure 6). This may be a generalisable outcome of DCA treatment of cancers that in our experiments correlated with the *in vivo* growth inhibitory efficacy of DCA. A reduced ability to synthesise the fatty acids required for membranes during proliferation would be expected to slow tumour growth rates, unless lipid scavenging pathways can compensate. The increase in malonic acid in DCA-treated 4T1s (Figure 5B) may point to either increased supply of this fatty acid precursor to assist fatty acid synthesis, or accumulation arising after 4T1 cells up-regulate lipid scavenging pathways. Further studies will be required to determine whether either of these adaptations contribute to the insensitivity of 4T1s to DCA.

While we do not know for sure the decrease in PC tail length is causative biology in terms of DCA growth inhibition, this result demonstrates how small changes in metabolic parameters and an overall effect may be detected by examining multiple points in the pathway, rather than only examining individual genes or metabolites. This may occur, even when no gene enrichment is highlighted by GO analysis, and when changes do not meet standard statistical cutoff criteria for significance. Further, a change in PC lipid profile can have anti-cancer impact beyond growth inhibition, as it can modify the physical properties of membrane lipid bilayers and can impact on processes such as autophagy and lipid signalling, which is implicated in apoptosis and drug resistance in cancer [97]. Further investigation is necessary to explore if these downstream metabolic effects of DCA can contribute to cancer therapy efficacy of other chemotherapy agents, even in the absence of growth inhibition.

### Conclusions

Thus, correlations of natural abundance isotopes with metabolic features suggest that δ^13^C and δ^15^N values are determined by multiple pathways in mouse mammary tumours and may correlate with DCA efficacy in vivo. The measurement of δ^15^N in these mouse mammary tumour models pointed to N-containing lipids as potential downstream mediators of DCA efficacy. Studies intervening in the lipid pathways are required to demonstrate causality for DCA, and studies with compound-specific isotope analysis would be necessary to provide direct evidence that intermediates of choline metabolism are ^15^N-depleted, or that polar heads of N-containing lipids are effectively ^15^N-depleted [43]. Further examination of δ^15^N for biomarker development in cancer is warranted.

## List of Abbreviations

BrCa: breast cancer
PDK: pyruvate dehydrogenase kinase
PDH: pyruvate dehydrogenase complex
TCA: tricarboxylic acid
DCA: dichloroacetate
IRMS: isotope ratio mass spectrometry
TNBC: triple negative breast cancer
HER2+: HER2 positive breast cancer
DE: differential expression

## Declarations

## Acknowledgements

The authors acknowledge the PLI of the CEISAM laboratory– ‘Isotope Platform in the Pays de la Loire’, as part of the “Infrastructure de recherche ligérienne” program launched by the Pays de la Loire region in collaboration with the European Commission, which supports research, development and innovation through the Plateforme Ligérienne d’Isotopie.

The authors thank Dr Ramon Sun for critically reading the manuscript, and the staff at the Central Facility for Genomics, Griffith University for their support with the NanoString gene expression analysis platform.

## Author contribution

Conceptualization, I.T, A.C.B, G.T; methodology, I.T, A.C.B, M.P.M.L, J.B, A-M.S, S.B, M.C; formal analysis, I.T, A.C.B, G.T; resources, A.C.B, G.T; data curation, I.T; writing-original draft preparation, I.T, A.C.B, G.T; writing-review and editing, I.T, A.C.B, M.P.M.L, J.B, A-M.S, S.B, M.C, G.T; supervision, I.T, A.C.B, G.T.

## Availability of data and materials

The dataset used and/or analyzed for this study is available upon reasonable request.

## Consent for publication

Not applicable

## Ethics approval

The study was approved by the Australian National University Animal Ethics Experimentation Committee (Protocols A2011-008 and A2014_19) under the guidelines established by the Australian National Health and Medical Research Committee.

