## Supplementary data for "Breast cancer metabolism and responsiveness to dichloroacetate: relationships with ^15^N and ^13^C natural abundance"

**Supplementary material**

**Table S1**. Common abbreviations used in figures for lipid molecular species.

| Abbreviation | Meaning |
|---|---|
| LPC | Lyso phosphatidyl-choline |
| PC | Phosphatidyl-choline |
| LPG | Lyso phosphatidyl-glycerol |
| PG | Phosphatidyl-glycerol |
| LPE | Lyso phosphatidyl-ethanolamine |
| PE | Phosphatidyl-ethanolamine |
| dMePE | Dimethyl phosphatidyl-ethanolamine |
| LPI | Lyso phosphatidyl-inositol |
| PI | Phosphatidyl-inositol |
| PL | Phospholipids |
| PS | Phosphatidyl-serine |
| DG | Diacylglycerol (diglycerides) |
| TG | Triacylglycerol (triglycerides) |
| PA | Phosphatidic acid |
| CL | Cardiolipin |
| Cer | Ceramide |
| CB | Cerebroside |
| diHexCB | Cerebroside with two glucosyl/galactosyl residues in polar head |
| monoHexCB | Cerebroside with one glucosyl/galactosyl residues in polar head |
| SM | Sphingomyelin |
| X(n:i, m:j) | X esterified with $C_n$ and $C_m$ fatty acid chains, with i and j unsaturations, respectively. Example, PC(18:3, 18:3): phosphatidyl-choline esterified with two $C_{18}$ chains carrying 3 unsaturations (linolenic type fatty acid). |

**Table S2.** Differentially expressed genes (unadjusted p<0.1) involved in metabolism from Nanostring Mouse Metabolism Panel on V14 mammary tumours, with and without DCA treatment.

| **Gene symbol** | **Panther protein class** | **GO Pathway** | **Gene name** |
|---|---|---|---|
| | **Enzyme activities / metabolism** | | |
| **Lipid metabolism** | | | |
| Pla2g15 | acyltransferase | OA | Phospholipase A2 group XV |
| Hacd2 | dehydratase | OA | Very-long-chain (3R)-3-hydroxyacyl-CoA dehydratase 2 |
| Acot12 | esterase | OA | Acetyl-coenzyme A thioesterase |
| **Amino acid / amine metabolism / anaplerosis** | | | |
| Odc1 | decarboxylase | OA | Ornithine decarboxylase |
| Ass1 | ligase | OA | Argininosuccinate synthase |
| Duox1 | oxidase | - | NAD(P)H oxidase (H2O2-forming) |
| Gpt | transaminase | SM | Alanine aminotransferase 1 |
| Got2 | transaminase | OA | Aspartate aminotransferase, mitochondrial |
| Idh1 | dehydrogenase | OA | Isocitrate dehydrogenase [NADP] cytoplasmic |
| **Other (nucleotides, TCA cycle etc)** | | | **Other (nucleotides, TCA cycle etc)** |
| Gmps | ligase | OA | GMP synthase [glutamine-hydrolyzing] |
| Ampd1 | deaminase | SM | AMP deaminase 1 |
| Inmt | methyltransferase | - | Indolethylamine N-methyltransferase |
| Adh7 | dehydrogenase | OA | All-trans-retinol dehydrogenase [NAD(+)] ADH7 |
| | **Other genes in GO SM pathway** | | |
| Rbp4 | transfer/carrier protein | SM | Retinol-binding protein 4 |
| Srebf2 | basic helix-loop-helix TF | SM | Sterol regulatory element-binding protein 2 |
| OA | Organic acid metabolic process / oxoacid metabolic process / carboxylic acid metabolic process | | |
| SM | Small molecule metabolic process | | |
| TF | Transcription factor | | |

**Table S3.** Non-metabolic, differentially expressed genes (unadjusted p<0.1) in V14 tumours, with and without DCA treatment.

| | | | |
|---|---|---|---|
| | **Other DE genes, not metabolic, not enriched in GO pathways** | | |
| | **Binding proteins etc** | | |
| Arpc4 | actin or actin-binding cytoskeletal protein | | Actin-related protein 2_3 complex subunit 4 |
| Npm1 | chaperone | | Nucleophosmin |
| Hspa4 | Hsp70 family chaperone | | Heat shock 70 kDa protein 4 |
| Wnt2 | intercellular signal molecule | | Protein Wnt-2 |
| Traf6 | scaffold/adaptor protein | | TNF receptor-associated factor 6 |
| Rbbp5 | signalling /binding protein | | Retinoblastoma-binding protein 5 |
| | **Transcription factors** | | |
| Hsf2 | winged helix/forkhead transcription factor | | Heat shock factor protein 2 |
| Zfp65 | C2H2 zinc finger transcription factor | | Zfp71-rs1 protein |
| | **Immune-related proteins** | | |
| Fpr1 | G-protein coupled receptor | | fMet-Leu-Phe receptor |
| Ccl4 | cytokine | | C-C motif chemokine 4 |
| Cpa3 | metalloprotease | | Mast cell carboxypeptidase A |
| Tpsb2 | serine protease | | Tryptase beta-2 |
| Tpsab1 | serine protease | | Tryptase |
| Ms4a1,Ms4a2 | transporter | | B-lymphocyte antigen CD20 (Ms4a1), High affinity immunoglobulin epsilon receptor subunit beta (Ms4a2) |

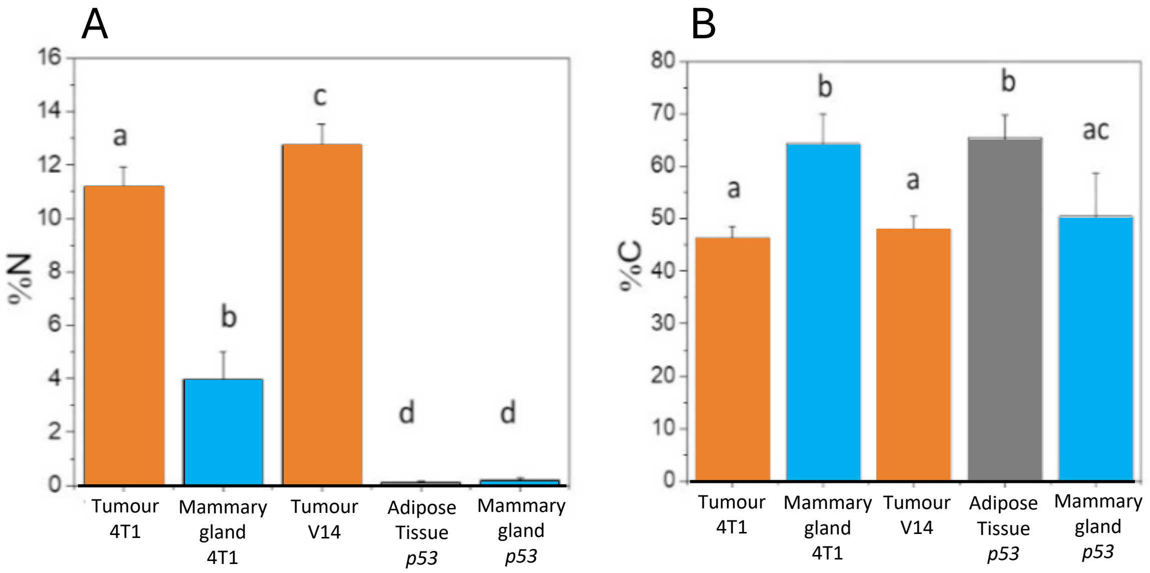

**Figure S1**. **Elemental content in tissues and tumours. (A)** N content (% of dry weight) of mammary gland, adipose tissue and tumour samples. V14 tumours had a higher %N compared to 4T1 tumours ($P < 0.01$), while both tumour types had far more N than mammary gland and adipose tissue. **(B)** C content (% of dry weight). Both 4T1 and V14 tumours had less carbon ($P < 0.05$) than mammary gland and adipose tissue. Lower case letters stand for statistical classes ($P < 0.05$).

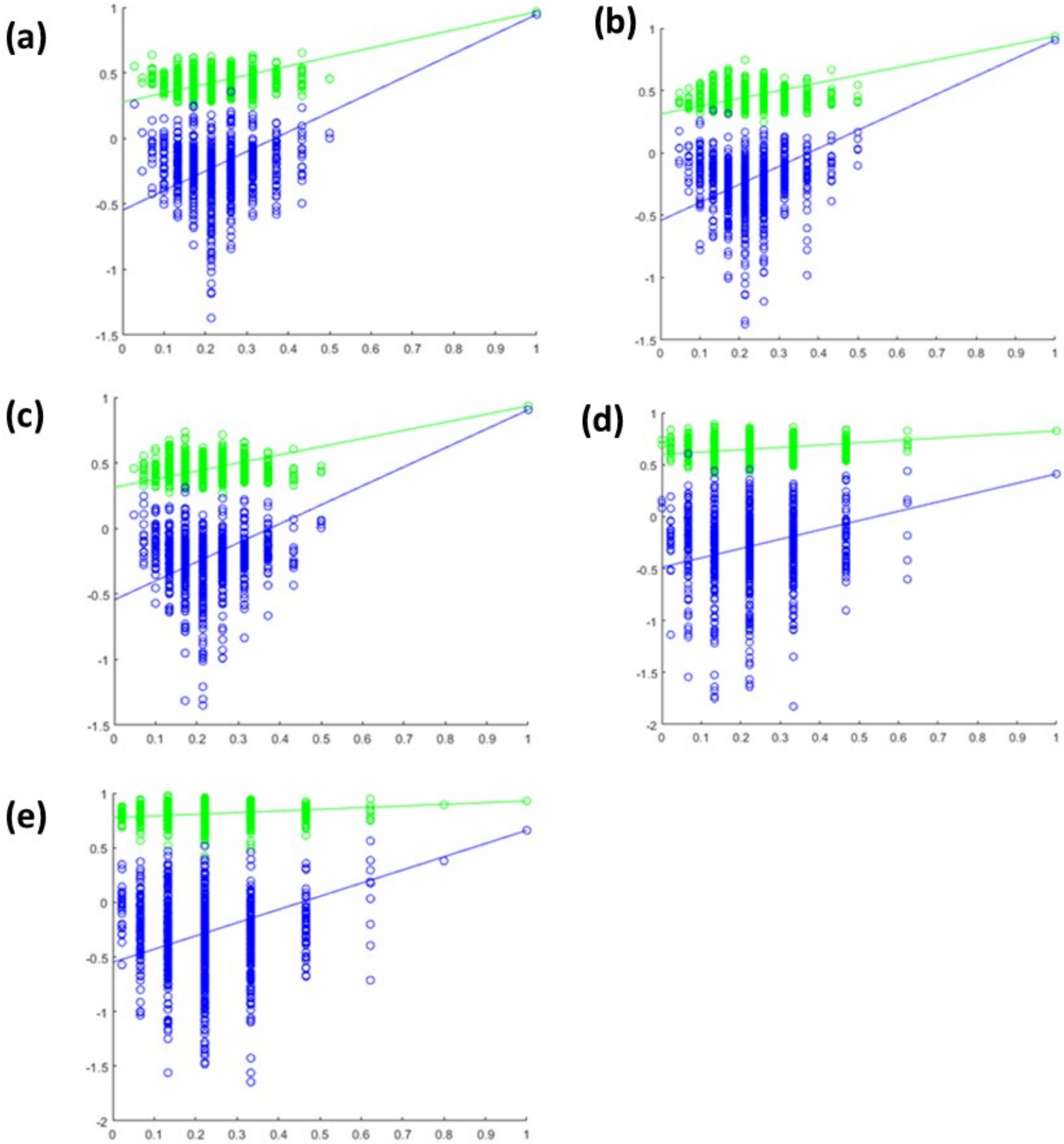

**Figure S2. Permutation tests obtained for: (a)** the C-OPLS-DA model between mammary tumours (n=21) and non-tumour samples (n=23); **(b) and (c)** the C-OPLS-R model on the same samples as (a) but with **(b)** $\Delta\delta^{15}N$ as the quantitative variable, or **(c)** $\Delta\delta^{13}C$ as the quantitative variable; **(d)** the C-OPLS-DA model between DCA-treated mammary tumours (n=10) and the untreated mammary tumours (n=11); and **(e)** the C-OPLS-R model on the same samples as (d) but with $\Delta\delta^{15}N$ as the quantitative variable. The permutation tests were performed with 1000 iterative rounds.

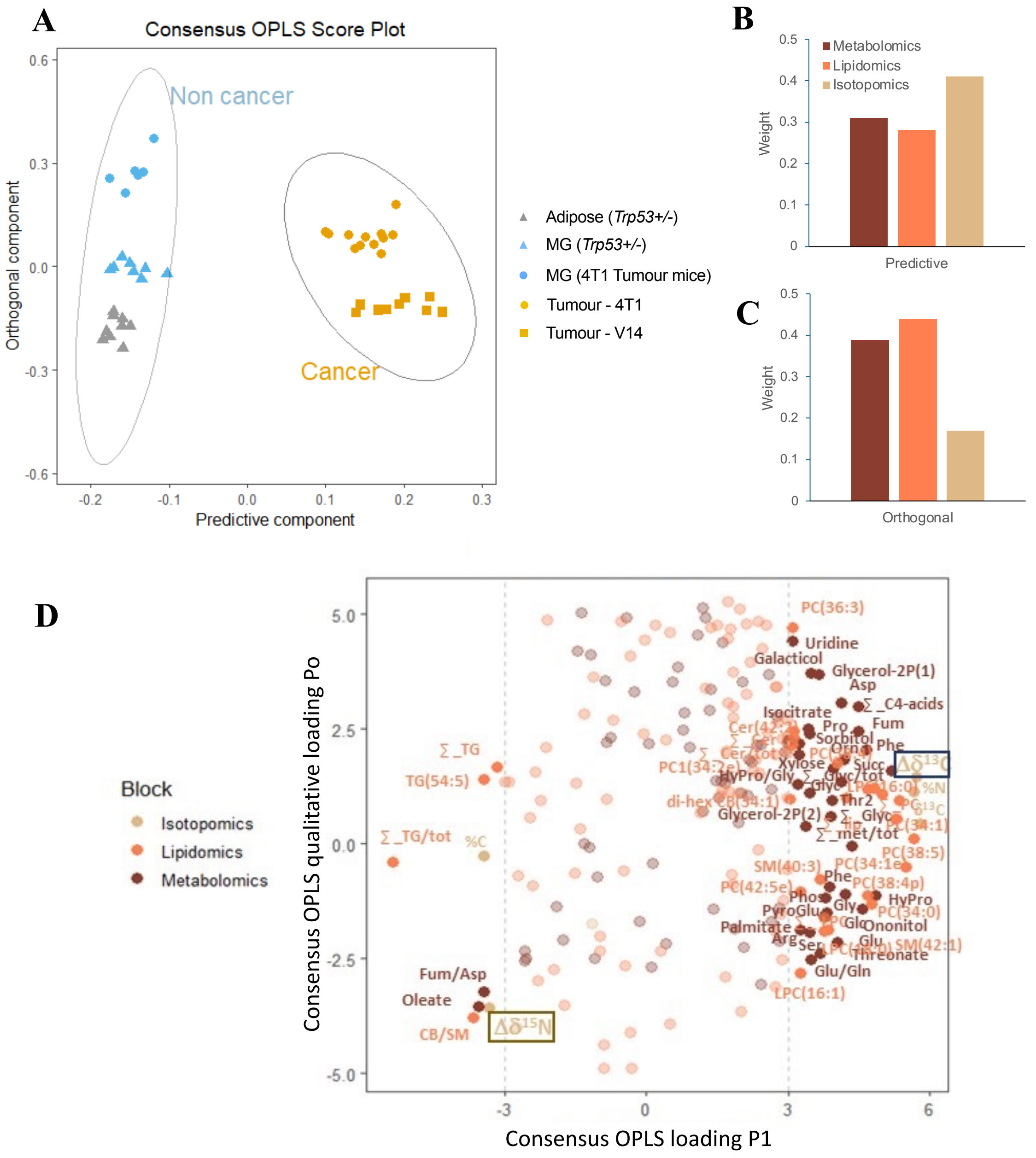

**Figure S3. Most significantly different features (combined metabolomics, lipidomics, and isotopic datasets) when comparing tumour and non-tumour tissues. (A)** Score plot of the C-OPLS-DA ($R^2Y = 0.97$ and $Q^2Y = 0.95$), which discriminates the tumour from the non-tumour tissues (predictive component). The ellipses are for a normal distribution with a level of 0.98. **(B)** and **(C)** Contributions of the metabolomic, lipidomic and isotopic blocks to **(B)** the predictive component and **(C)** the orthogonal component of the score plot in (A). **(D)** The C-OPLS loading plot highlights the significant features distinguishing between tumour and non-tumour tissues among isotopomic, metabolomic and lipidomic datasets. The features are colour-coded according to which block they belong. The best features (loading value below – 3 or above +3 for P1) were annotated (abbreviations in Table S1).

**Supplementary details of the statistical analyses:**
Additional feature details of the datasets analysed (isotopomic, metabolomic and lipidomic) and statistical analysis performed are provided here for each relevant figure of the manuscript.

**Statistical Analysis - Figure S3.**
A combined (multiblock) analysis of the three datasets, isotopic (7 features), metabolomic (96 features) and lipidomic (112 features), was conducted. Autoscaling was applied to datasets and C-OPLS discriminant analysis was carried out with samples grouped for comparison of tumours ($n$ = 21) with non-tumour samples ($n$ = 23), regardless of the tumour type or mouse treatment / genotype group. The model was found to be optimal with two latent variables (one predictive and one orthogonal) using leave-one-out cross-validation, leading to high model fit ($R^2 = 0.97$) and prediction performance ($Q^2 = 0.95$). Further confirmation of model robustness was carried out using permutation tests (1,000 iterations, Figure S2a).

**Statistical Analysis - Figure 2.**
Potential links between metabolic features and isotope abundance was explored via a covariation analysis between $\Delta\delta$ values and metabolomic and lipidomic feature peak areas. That is, two additional C-OPLS regression models were computed using $\Delta\delta^{15}N$ or $\Delta\delta^{13}C$ as Y response variables and feature levels as the X axis. Cross-validation was carried out to assess model optimal complexity and performance. Three latent variables (1 predictive + 2 orthogonal) were generated in the $\Delta\delta^{15}N$ model (with $R^2 = 0.71$ and $Q^2 = 0.55$) while four components (1 predictive + 3 orthogonal) were generated in the $\Delta\delta^{13}C$ model (with $R^2 = 0.99$ and $Q^2 = 0.96$). Model robustness was assessed using permutation tests (1,000 iterations), as shown in Figure S2b and S2c. Volcano plots were used to identify the best potential drivers of $\Delta\delta^{15}N$ and $\Delta\delta^{13}C$ (Fig. 2) in the tumour and non-tumour tissues.

**Statistical Analysis - Figure 3.**
To identify metabolic features that contribute to the cellular content of $^{15}N$ in these tumours, C-OPLS regression analysis was undertaken on tumour samples only, with $\Delta\delta^{15}N$ as the Y quantitative response variable. Monte Carlo uninformative variable elimination MCUVE-PLS was applied with two latent variables and 1,000 re-samplings for uninformative feature elimination. The resulting multiblock dataset comprised 50 features from the metabolite and lipid datasets. C-OPLS analysis with cross-validation generated seven latent variables (1 predictive + 6 orthogonal) with satisfactory performance ($R^2 = 1.00$, $Q^2 = 0.39$; outputs of permutation tests are provided in Figure S2e). Since 4T1 and V14 tumours had different average $\Delta\delta^{15}N$ values (Figure 1C and Figure 3B), centering and unit-variance scaling was done to eliminate the tumour model effect. A volcano plot (Figure 3D) was constructed combining C-OPLS loadings ($p_{corr}$) with $P$-values computed from linear regression (after Benjamini-Hochberg false discovery rate correction) between features and $\Delta\delta$.

**Statistical Analysis - Figure 4.**
A multivariate analysis was carried out to explore the effect of DCA on metabolism, using 4T1 and V14 tumours treated and untreated with DCA (i.e. four sample groups). At first, the C-

OPLS-DA model showed poor performance, and therefore a mid-level data integration analysis was implemented. To do so, Montecarlo uninformative variable elimination (UVE) was done separately on the three datasets (blocks) in order to select the most reliable features. Autoscaling was then applied to each dataset. Two components were found to be optimal to apply MCUVE-PLS, with 1,000 Monte Carlo resampling rounds. The resulting (trimmed) combined dataset comprised 1 isotopic feature, 47 metabolites and 58 lipid features. C-OPLS-DA analysis was then conducted with the trimmed dataset (score plot shown in Figure 4A). Two latent variables (1 predictive + 1 orthogonal) were found to be optimal using leave-one-out cross-validation, leading to satisfactory model fit ($R^2 = 0.83$) and moderate prediction performance ($Q^2 = 0.41$). Permutation tests were then performed (1,000 iterations), showing acceptable results (Figure S2D).
